# CIP2A is a biomarker for progression and poor prognosis in cutaneous squamous cell carcinoma

**DOI:** 10.64898/2026.11.19.767767

**Authors:** Liisa Nissinen, Panyi Tran, Lauri Heiskanen, Jutta Huvila, Jaakko S. Knuutila, Jukka Westermarck, Veli-Matti Kähäri, Pilvi Riihilä

## Abstract

The incidence of cutaneous squamous cell carcinoma (cSCC) is rising worldwide, highlighting the need for novel therapeutic options for recurrent and metastatic disease. Protein phosphatase 2A (PP2A) is a keyt tumor suppressor, and its inhibition promotes malignant progression. Here, we investigated the role of the cancerous inhibitor of PP2A (CIP2A) in cSCC progression. RNA-sequencing, oligonucleotide array-based expression, and qRT-PCR analysis demonstrated elevated CIP2A mRNA levels in cSCC cells compared with normal human epidermal keratinocytes (NHEKs). Western blotting confirmed higher CIP2A protein production in cSCC cells than in NHEKs. Analysis of single-cell RNA-sequencing data from cSCC showed CIP2A expression in epithelial cells and fibroblasts. Silencing CIP2A with specific siRNAs reduced cSCC cell growth and viability by inhibiting MEK1 and ERK1/2 phosphorylation. *In vivo*, strong CIP2A immunostaining was observed in tumor cells of primary cSCCs and cSCC metastases, whereas staining was absent or weak in all actinic keratoses and most cSCC *in situ* samples. Kaplan-Meier analysis revealed that patients with high CIP2A expression had significantly worse disease-specific survival than those with low expression. These findings identify CIP2A as a clinically relevant oncogene in cSCC progression, with elevated expression in invasive primary tumors and metastases. CIP2A may serve as a biomarker of cSCC progression and metastasis, as well as a potential therapeutic target in advanced and metastatic cSCC.

## INTRODUCTION

Cutaneous squamous cell carcinoma (cSCC) arises from epidermal keratinocytes and is the most common metastatic skin cancer, with an incidence that is increasing worldwide (Nehal and Bichakjian 2018). The metastasis rate of primary cSCC is estimated at 3–5%, and outcomes for patients with metastatic disease remain poor: over 70% of individuals with metastatic cSCC (mcSCC) die within three years (Knuutila et al 2020; Nagarajan et al 2019). Overall, cSCC is estimated to account for approximately 20% of skin cancer–related mortality (Nehal and Bichakjian 2018). Actinic keratosis (AK), a ultraviolet (UV) radiation-induced premalignant lesion, can progress to malignant cSCC *in situ* (cSCCIS, Bowen’s disease) and subsequently to invasive and mcSCC (Ratushny et al 2012). Long-term exposure to UV radiation is the primary risk factor for cSCC development, while additional risk factors for cSCC initiation and progression include chronic inflammation, chronic dermal ulcers, and immunosuppression (Ratushny et al 2012).

Protein phosphatase 2A (PP2A) is a key serine/threonine phosphatase that plays a critical role in counteracting the hyperactive kinases characteristic of many malignancies. It has tumor suppressor functions and accounts for the majority of cellular serine/threonine phosphatase activity (Sablina et al 2010). Inactivation of PP2A is a critical requirement for the transformation of human cells (Sablina et al 2010). Several endogenous PP2A inhibitors have been identified, including cancerous inhibitor of PP2A (CIP2A), protein phosphatase methylesterase-1 (PME-1) and SET. These inhibitors of PP2A are frequently upregulated in cancer, with expression levels varying across malignancies and among subtypes of individual cancers (Junttila et al 2007; Puustinen et al 2009; Westermarck and Hahn 2008). Inhibition of PP2A activity by these endogenous inhibitors promotes oncogenic kinase activation and transcription factor signaling, thereby facilitating tumor development (Kauko et al 2020).

Among these inibitors, CIP2A is the most prevalent oncoprotein. Its overexpression has been reported in a broad spectrum of cancers, including head and neck squamous cell carcinoma (HNSCC), as well as breast, colon and gastric cancers, where it is associated with poor prognosis (Junttila et al 2007; Côme et al 2009; Khanna et al 2009). High CIP2A expression has also been linked to poor therapeutic response in several cancers (Khanna and Pimanda 2016). Mechanistically, CIP2A promotes MYC protein stability in human cancer cells (Junttila et al 2007). It also interacts with the DNA repair scaffold protein TopBP1 to coordinate DNA damage–induced mitotic checkpoint control and proliferation, thereby driving the initiation and progression of basal-like breast cancers (Laine et al 2021). In addition, CIP2A has been shown to enhance cellular proliferation and tumor growth *in vivo* in human HNSCC (Junttila et al 2007). However, whether CIP2A expression can be used to estimate the aggressiveness of cSCC of skin remains largely unexplored.

In this study we have investigated the role of CIP2A in the progression of cSCC. Our results show that CIP2A expression is elevated in cSCC tumor cells compared with normal epidermal keratinocytes (NHEKs) in culture, whereas SET and PME-1 are not significant regulated. The results also demonstrate, that CIP2A regulates the growth and viability of cSCC cells by modulating the activity of MEK1-ERK1/2 signaling pathway. *In vivo,* CIP2A is strongly overexpressed in tumor cells of invasive cSCCs and metastases, and elevated CIP2A expression correlates with poor disease-specific survival. Together, these findings identify CIP2A as a biomarker and a potential therapeutic target in locally advanced and metastatic cSCCs.

## MATERIALS AND METHODS

### Ethical issues

Approval for tissue specimens and the collection of tissue samples was obtained from the Ethics Committee of the Hospital District of Southwest Finland, Turku, Finland (187/2006) and Auria Biobank’s Scientific Steering Committee (AB15-9721). The study was performed in accordance with the ethical guidelines of the Declaration of Helsinki. Each patient gave their written, informed biobank consent. Registry study approval for collection, and use of clinical and histopathological data was obtained from the Turku University Hospital Clinical Research Centre (TO5/042/18).

### Human tissue samples and immunohistochemical analysis

Human tissue samples were obtained from Auria Biobank and the archives of Turku University Hospital. Tissue samples were originally obtained during normal diagnostic or therapeutic procedure. Human tissue used for immunohistochemical (IHC) analysis consisted of formalin-fixed, paraffin embedded (FFPE) samples as follows: normal skin (n=67), AK (n=73), cSCCIS (n=59), cSCC (n=162) and cSCC metastasis (n=63). As cSCC cell culture data showed increase in CIP2A expression, but not in other PP2A inhibitors (PME-1 and SET), CIP2A immunostainings were performed in patient samples. IHC was performed on tissue microarrays (TMA) blocks using rabbit polyclonal CIP2A antibody (HPA039570, Sigma-Aldrich, St. Louis, MO). For semiquantitative evaluation, CIP2A immunostaining was scored as follows: negative/weak staining (-) and strong staining (+). Semiquantitative evaluation was performed independently with digital microscopy by two observers (P.T. and P.R.) and confirmed with pathologist (J.H.). For negative control stainings the primary antibody was omitted and replaced by PBS.

### Cell culture

Human primary non-metastatic (UT-SCC-12A, UT-SCC-91, UT-SCC-105, UT-SCC-111, UT-SCC-118) and metastatic (UT-SCC-7, UT-SCC-115, UT-SCC-59A) cSCC cell lines were established from surgically removed cSCCs and cultured as previously described (Riihilä et al 2014). The authenticity of all cSCC cell lines has been verified by short tandem repeat profiling (DDC Medical, Fairfield, OH) (Farshchian et al 2017). Keratinocytes, established from skin samples obtained from patients undergoing surgery for mammoplasty at Turku University Hospital (n=7, Turku, Finland) and from PromoCell (n=1, Heidelberg, Germany) were cultured as previously described (Riihilä et al 2017; Farshchian et al 2017).

### Tissue RNA

Normal human skin samples (n = 6) were obtained from the Turku University Hospital (Turku, Finland) from the upper arm of healthy volunteers or during mammoplasty surgery. Human primary cSCC samples (n = 6), removed during surgery, were also obtained from Turku University Hospital (Riihilä et al 2020). Total RNA was isolated as previously described (Riihilä et al 2020; Viiklepp et al 2025).

### RNA extraction

Extraction of RNA was performed as described previously (Rahmati Nezhad et al 2022; Riihilä et al 2020). For the whole transcriptome sequencing total RNA was isolated using miRNAeasy Mini kit (Qiagen, Chatworth, CA) according to the manufacturer’s protocol.

### Whole transcriptome and oligonucleotide array-based expression analysis

RNA sequencing analysis was performed using Illumina HiSeq2500 system using paired-end sequencing chemistry with 100 bp read length (Illumina, San Diego, CA) at the Finnish Functional Genomics Centre, University of Turku, Åbo Akademi University and Biocenter Finland as described previously (Nissinen et al 2024). Affymetrix human U133 Plus 2.0 gene chip was used for gene expression profiling (Finnish Microarray and Sequencing Centre, Turku Center for Biotechnology, Turku, Finland). Normalization of the data was prepared with RMA (Chipster, CSC, Finland). The microarray data (accession number GSE66368) and RNA sequencing data (accession number GSE252347) are available at the GEO (Gene Expression Omnibus, NCBI; http://www.ncbi.nlm.nih.gov/geo/) (Riihilä et al, 2015).

### Quantitative real-time PCR

Preparation of cDNA was done as described previously (Riihilä et al, 2015). Quantitative real-time PCR (qRT-PCR) was performed to determine the levels of CIP2A and PPME1 (PME-1) (Mäkelä et al 2019) and SET Assay on demand (4331348SET, Thermo Fisher Scientific (Waltham, MA, USA) to determine the levels of SET. Glyceraldehyde-3-phosphate dehydrogenase and β-actin were used as housekeeping controls (Farshchian et al 2011, Riihilä et al 2015). qRT-PCR was performed using Applied Biosystems 7900HT Fast Real-Time PCR System.

### Western blot analysis

Proteins were analyzed after fractionation by 10% SDS-PAGE. Antibodies against CIP2A (sc-80659), PME-1 (sc-20086) and SET (sc-133138) (all from Santa Cruz Biotechnology, Santa Cruz, CA) were used. ERK1/2 and MEK1 were detected with antibodies specific for phosphorylated forms and total proteins (#9101, #9154, #9102, #9124, respectively; all from Cell Signaling Technology, Danvers, MA, US). As loading control antibody against ß-actin (Sigma-Aldrich) was used. Fluorescently-labeled secondary antibodies (LI-COR Biosciences, Lincold, NE) and the LI-COR Odyssey CLx imaging system was used to quantitate the protein expression.

### Single-cell RNA-sequencing data analysis

The single-cell RNA-sequencing datasets were obtained from Gene Expression Omnibus database. The dataset GSE193304 (Zou et al., 2023) was normalized by Bioconductor package Seurat (version 5) (Hao et al, 2024) implemented in Chipster (Kallio et al, 2011) using “LogNormalize” method and scaling factor of 10,000. The dataset was converted to expression matrix files by Seurat (version 5) and the expression matrix was normalized as described earlier and clustered using Seurat, version 5, implemented in Chipster (Kallio et al, 2011). Annotation was done in Chipster using SingleR (Aran et al, 2019) and human primary cell atlas as CellDex reference. The percentage of cells expressing a specific marker and the average expression of that marker in each cluster were extracted from Seurat object using Chipster.

### CIP2A knock-down with small interfering RNAs

Small interfering RNAs (siRNA) for CIP2A (75 nM) were described previously (Laine et al 2021). Nonspecific siRNA (Qiagen, India) was used as a negative control. Cells were transfected using siLentFect^TM^ Lipid Reagent (Bio-Rad Laboratories, Hercules, CA) (Riihilä et al 2015).

### Cell growth assay

Cells were seeded (7.5 × 10^3^ cells/well) on 96-well plates. The IncuCyte S3 real-time cell imaging system (Essen BioScience, Ann Arbor, MI) was used to study the growth of cSCC cells. Images were taken every 4 h by IncuCyte S3, and the relative confluence was analyzed by the instrument. Experiments were carried out with 5–8 parallel wells at every time point with three cSCC cell lines (UT-SCC-12A, -7 and -59A) as previously described (Nissinen et al 2024).

### Cell viability assay

To study the effect of CIP2A on cSCC proliferation, cells were transfected with negative control siRNA or CIP2A siRNAs (75 nM). Cell Counting Kit-8 (CCK-8, Dojindo, Japan) was used to measure cellular viability 24, 48, and 72 h after transfection according to manufacturer’s instructions (Riihilä et al 2020). The experiment was performed with 3-8 parallel wells in each time point with four cSCC cell lines (UT-SCC-12A, -91, -7 and -59A).

### Determination of the cell number

To study the effect of CIP2A on cell number, cells were transfected with negative control siRNA or CIP2A siRNAs (75 nM). SYTO 60 (Thermo Fisher Scientific, Waltham, MA) DNA staining method was used to determine cSCC cell number 24, 48, and 72 h after transfection. Briefly, cells were fixed with 4% paraformaldehyde and permeabilized by 0.2% Triton X-100. After that the cells were treated with SYTO 60 reagent and analyzed with LI-COR Odyssey CLx imaging system. The experiment was performed with 6-8 parallel wells in each time point with two cSCC cell lines (UT-SCC-12A and -91).

### Survival analysis

Patients were stratified into two groups according to CIP2A expression one group represented the high CIP2A expression and other group represented the low or negative expression whether the tumor were metastatic or non-metastatic, and survival times were calculated from tumor detection to death, or the end of 5-year follow up period. Overall survival and disease-specific survival analyses were implemented.

### Statistical analysis

Two independent samples T-test was used for the cell growth and viability assays and analysis of cell number. Two-tailed Mann-Whitney U test was used for qRT-PCR. Chi-square test was used for IHC analysis. The Kaplan–Meier method with log-rank test was applied to generate survival curves and define survival estimates. JMP (2025 JMP® Version 18, Student Edition. JMP Statistical Discovery LLC, Cary, NC.) was used to perform the survival analyses. P values < 0.05 were considered statistically significant.

## RESULTS

### Expression of CIP2A is upregulated in cSCC cells

PP2A inhibitor expression profiles were determined in primary (n=5) and metastatic (n=3) cSCC cell lines, as well as in primary human epidermal keratinocytes (NHEKs) (n=5) from healthy skin, using oligonucleotide array–based expression profiling (Figure 1A) and RNA sequencing (NHEK, n=4) analysis (Figure 1B). Expression of CIP2A mRNA was upregulated in cSCC cells compared to NHEKs, whereas lower expression was detected in NHEKs (Figure 1A, B). The mRNAs for SET and PPME1 (PME-1) were expressed both in cSCC cells and NHEKs (Figure 1A, B). The expression of CIP2A, SET and PPME1 in NHEKs and cSCC cell lines was also determined with qRT-PCR (Figure 1C). Expression of CIP2A mRNA was significantly upregulated in cSCC cells compared with NHEKs, while no difference was noted in SET and PPME1 mRNA levels (Figure 1C). Western blot analysis of protein levels of CIP2A, SET and PME-1 showed, that CIP2A expression was low in NHEKs and upregulated in cSCC cells (Figure 1D). Expression of SET and PME-1 was detected both in cSCC cells and NHEKs (Figure 1D). Elevated mRNA levels of CIP2A were also detected in *in vivo* cSCC tumor tissue (n=6) compared with normal skin (n=6) (Figure 1E), while SET and PPME1 mRNA levels were lower in cSCC tumors compared to normal skin *in vivo* (Figure 1E).

**Figure 1.**
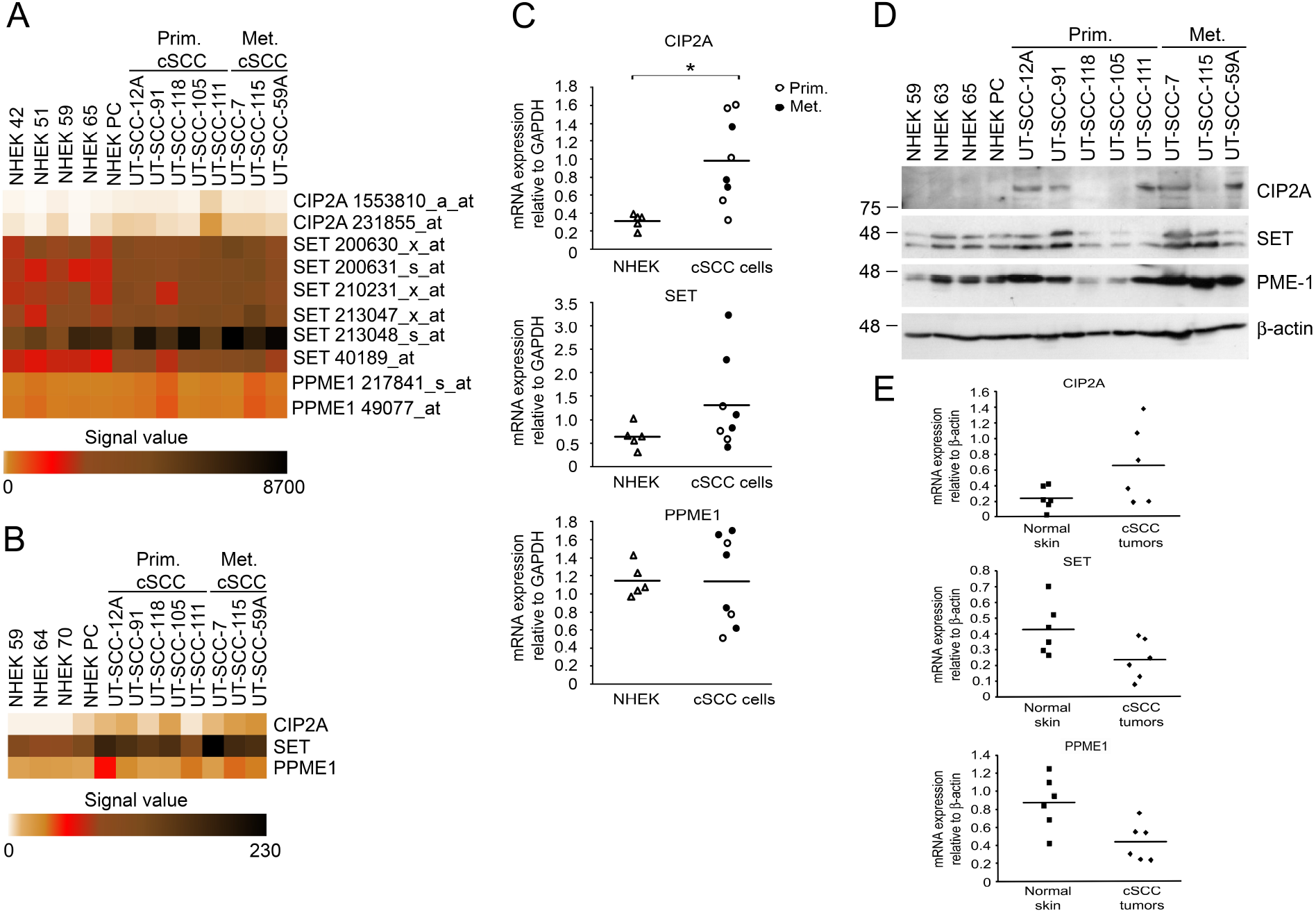
Expression of PP2A inhibitors in cSCC. (A) RNAs from normal human epidermal keratinocytes (NHEK, n=5), and primary (Prim. cSCC, n=5) and metastatic (Met. cSCC, n=3) cutaneous SCC cell lines were analyzed by Affymetrix gene chip technology. (B) The whole transcriptome sequencing analysis was performed with RNAs from NHEKs (n=4), and primary (n=5) and metastatic (n=3) cSCC cell lines. The normalized signal values are shown in heatmaps. (C) The mRNA levels of CIP2A, SET and PPME1 (PME-1) in NHEKs (n=5) and cSCC cell lines (n=8) were determined with qRT-PCR. (D) Production of CIP2A, SET and PME-1 proteins was analyzed by western blotting. Migration positions of molecular weight markers (in kDa) are shown on the left. (E) CIP2A, SET and PPME1 (PME-1) mRNA levels were determined from normal skin (n=6) and cSCC tumors (n=6) with qRT-PCR. Horizontal bars indicate mean for each group. Two-tailed Mann-Whitney U test. *p< 0.05.

Analysis of single-cell RNA-seq data from cSCC (Zou et al 2023) revealed CIP2A mRNA expression predominantly in epithelial cells (cluster 10) and fibroblasts (cluster 20) (Figure 2A and 2B). In contrast, SET and PPME1 were broadly expressed across multiple clusters (Figure 2A and 2B). Similarly, analysis of normal skin single-cell RNA-seq data (Zou et al 2023) showed CIP2A mRNA expression in epithelial cells (Figure 2C and 2D), whereas SET and PPME1 were widely expressed across several clusters (Figure 2C and 2D).

**Figure 2.**
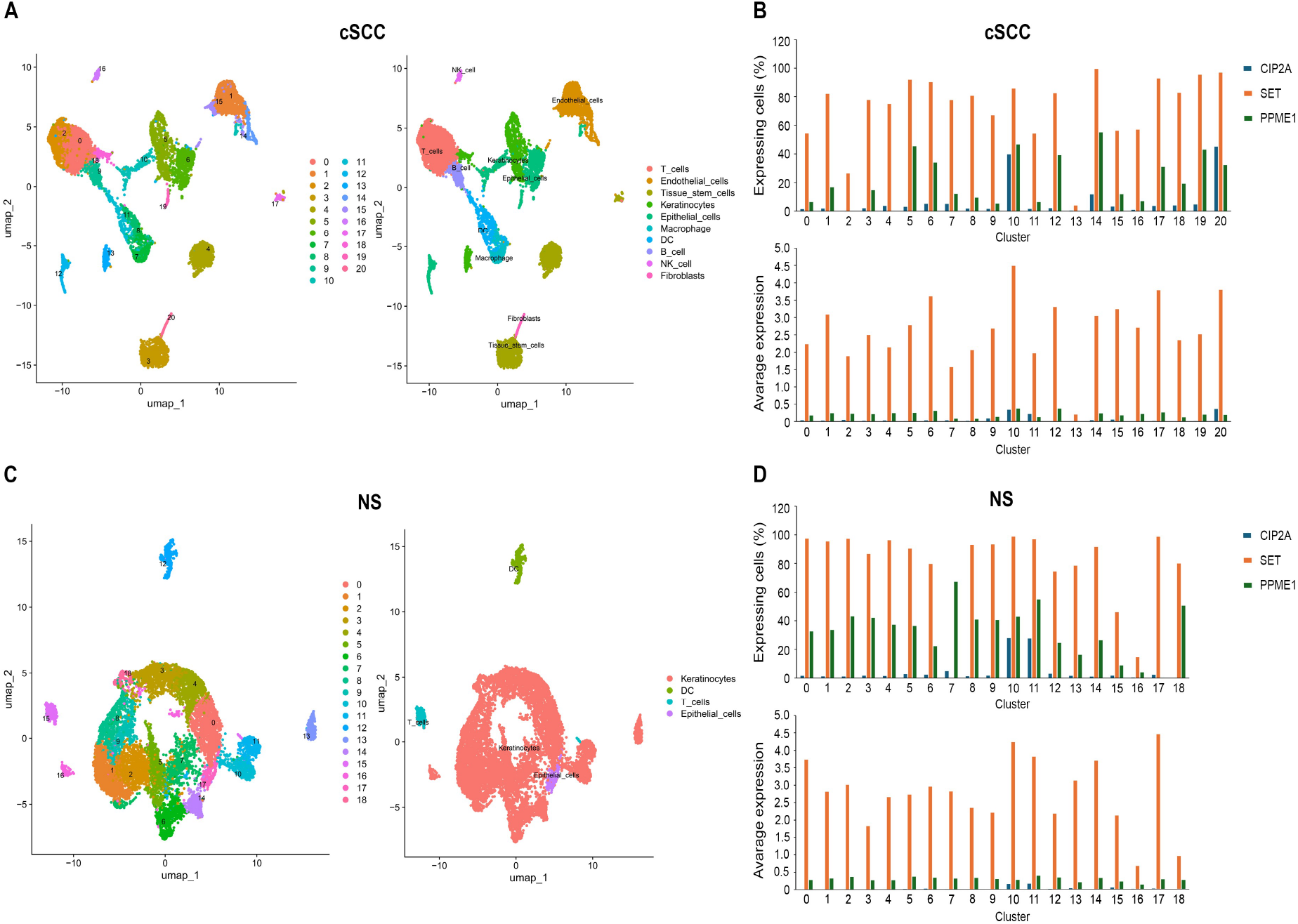
Expression of CIP2A, SET and PPME1 in cSCC *in vivo*. Single-cell RNA-sequencing dataset of cSCC (A,B) GSM5788502 and normal skin (NS) (C,D) GSM5788503 from Gene Expression Omnibus database was converted to expression matrix files by Seurat, version 5. The expression matrix of the tumor sample with the highest number of analyzed cells was normalized and clustered using Seurat, version 5, implemented in Chipster. Annotation was performed in Chipster using SingleR and human primary cell atlas as CellDex reference. (B,D) Upper panel: percentage of cells expressing mRNAs for CIP2A, SET and PPME1 in each cluster. Lower panel: average expression of the same genes in each cluster.

### CIP2A regulates cSCC cell growth and viability

Since CIP2A was the only PP2A inhibitor studied that showed weak or no expression in NHEKs but was highly expressed in cSCC, we examined its role by knocking down CIP2A expression with specific siRNAs (75 nM) (Figure 3A, Supplementary Figure S1). CIP2A knockdown significantly inhibited cSCC cell growth (Figure 3B, Supplementary Figure S2) and viability (Figure 3C, Supplementary Figure S3) of cSCC cells. Furthermore, CIP2A knockdown significantly reduced the number of cSCC cells (Figure 3D, Supplementary Figure S4).

**Figure 3.**
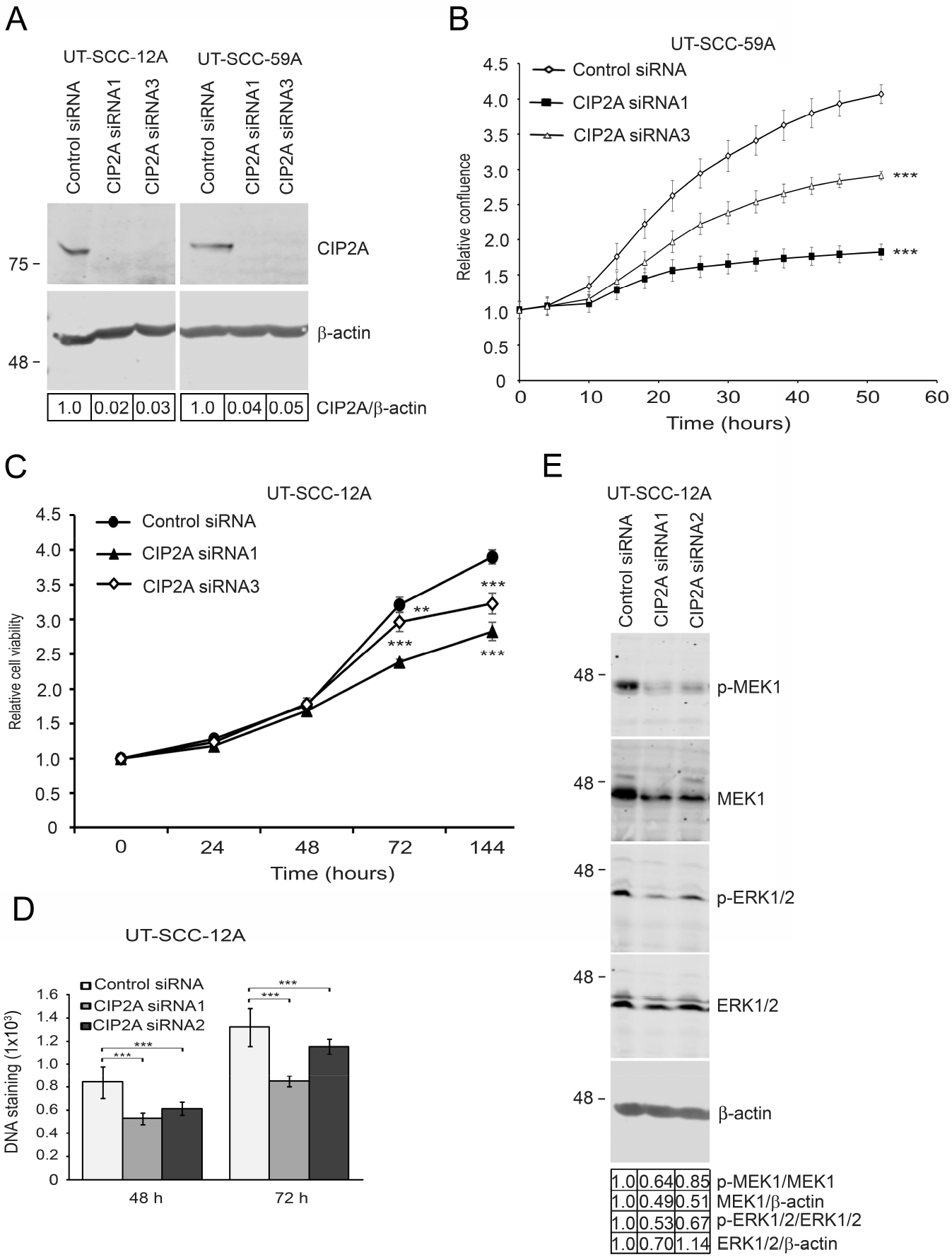
CIP2A regulates the growth of cSCC cells. Cutaneous SCC cells were transfected with 75 nM negative control (Control siRNA) or CIP2A siRNAs. (A) CIP2A level was determined with western blotting after 72 h. β-actin was used as a loading control. Migration positions of molecular weight markers (in kDa) are shown on the left. (B) The confluency of cSCC cells after CIP2A siRNA transfections was determined using IncuCyte S3 (n=8). (C) The number of viable cSCC cells was determined after siRNA transfection using CCK-8 assay at timepoints indicated (n=7-8). (D) DNA staining was performed 24, 48 and 72 h after siRNA transfection with SYTO 60 reagent and analyzed with LI-COR Odyssey CLx imaging system (n=8). (E) The levels of p-MEK1, p-ERK1/2, total MEK1 and total ERK1/2 in cell lysates of CIP2A siRNA transfected cSCC cells were determined by western blotting 72 hours after transfection. β-actin was used as a sample control. Migration positions of molecular weight markers (in kDa) are shown on the left. Two-tailed Student’s t-test. **p < 0.01, ***p < 0.001.

### CIP2A regulates phosphorylation of MEK1 and ERK1/2

Given the observed effects CIP2A silencing on cSCC cell growth, we next investigated the activity of key components of the MEK1–ERK1/2 signaling pathway, which plays an important role in cSCC development. As shown in Figure 3E, CIP2A knockdown inhibited the phosphorylation of MEK1 and ERK1/2 in cSCC cells.

### Increased expression of CIP2A in cSCC in vivo

To investigate the role of CIP2A in progression of cSCC *in vivo*, we analyzed its expression by IHC in TMAs comprising solar UV induced premalignant skin lesions (AK, n=73), cSCC in situ (SCCIS, n=59), invasive cSCC (n=162) and cSCC metastases (n=63). Prominent CIP2A expression was detected in tumor cells of primary cSCC tissue samples (Figure 4A) and metastases (Figure 4B). CIP2A expression was also noted in fibroblast-like cells within the tumor microenvironment in cSCC (Figure 4A). In contrast, only weak staining was detected in epidermal keratinocytes of normal skin (Figure 4C), AK (Figure 4D) and cSCCIS (Figure 4E).

**Figure 4.**
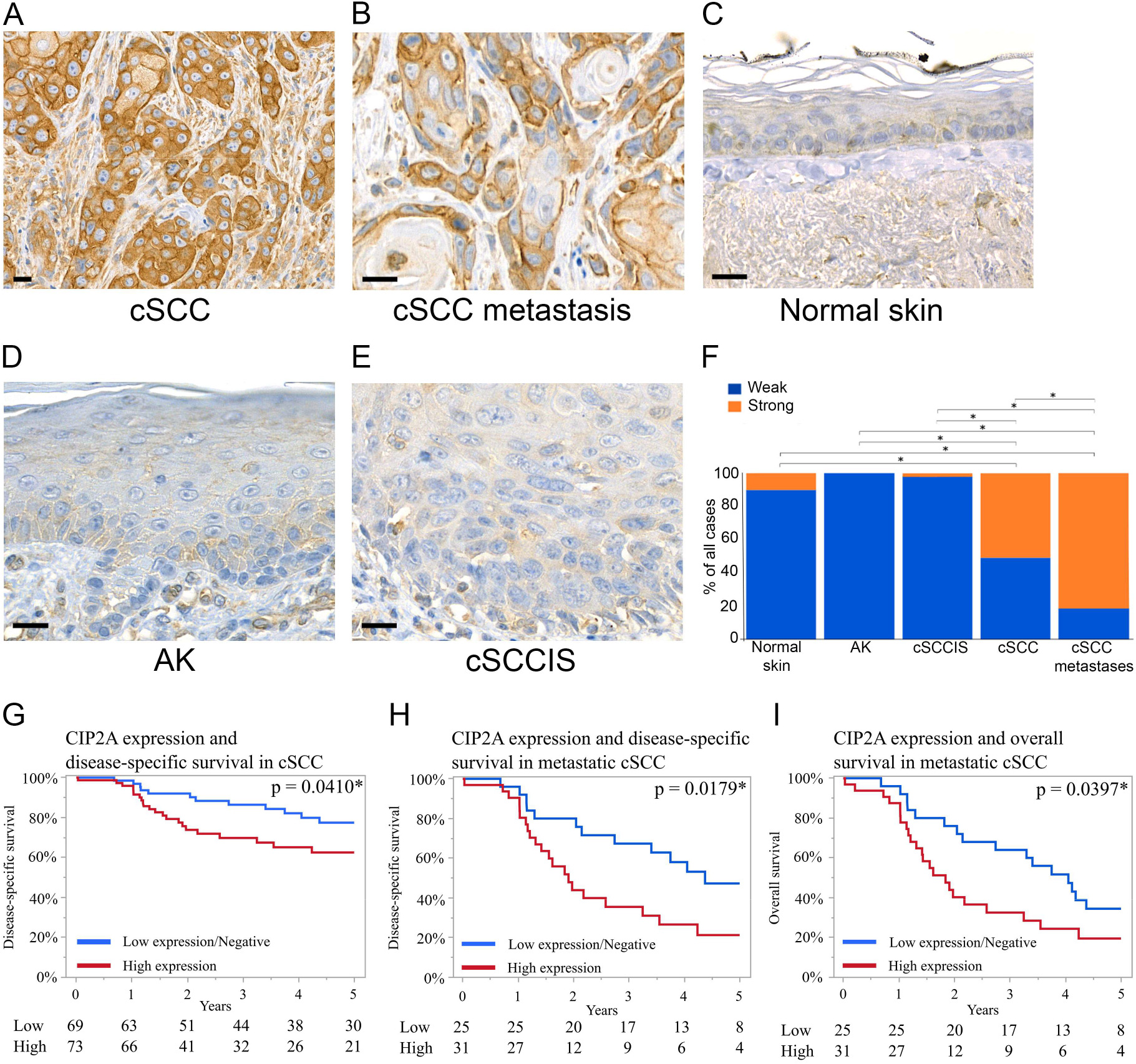
Increased expression of CIP2A is associated with poor prognosis in cSCC. (A–F) Immunohistochemical staining with a CIP2A antibody was performed on tissue samples from normal skin (n = 67), actinic keratosis (AK; n = 73), cSCC *in situ* (cSCCIS; n = 59), invasive cSCC (cSCC; n = 162 individual tumors) and cSCC metastases (n = 63 individual tumors). Representative stainings from each group are shown. (A, B) Increased CIP2A staining was observed in cSCC cells. (A-F) The CIP2A staining was stronger on the tumor cells in invasive cSCC and cSCC metastases compared to normal skin, AK and cSCCIS. (F) CIP2A immunostaining intensity was scored as weak or strong based on specific staining on the epithelial cells. The CIP2A staining in cells was significantly stronger in invasive cSCC and particularly in metastases compared with normal skin, AK, and cSCCIS. ∗P<0.0001, Chi-square test. Scale bars 50 μm. (G-I) Kaplan-Meier survival curves comparing high versus low CIP2A expression in patients with cSCC. The difference between groups was evaluated using the log-rank test. Survival time was calculated from time of tumor detection to death or the end of five-year follow up. Numbers at risk shown below each corresponding panel. (G) Disease-specific survival of all cSCC patients with high CIP2A expression compared to those with low expression (log-rank p = 0.0410). (H) Disease-specific survival of patients with mcSCC tumor with high and low CIP2A expression analyzed alone without patients with non-metastatic tumor (log-rank = 0.0179). (I) Overall survival of patients with mcSCC with high and low CIP2A expression (log-rank p = 0.0397).

Semiquantitative analysis revealed that CIP2A was significantly upregulated in cSCC tumor cells: 82 of 162 cases (51%) displayed strong CIP2A staining, whereas 80 of 162 cases (49%) were negative or weak (Figure 4F, Supplementary table S1). Tumor cells in metastases had the highest CIP2A expression with 51 of 63 cases (81%) showing strong staining and 12 of 63 cases (19%) being negative or weak (Figure 4F, Supplementary table S1). In AKs, CIP2A staining was negative or weak in all 73 cases (Figure 4F, Supplementary table S1). In cSCCIS, CIP2A staining was negative or weak in 58 of 59 cases (98%), with only 1 of 59 cases (2%) showing strong staining (Figure 4F, Supplementary table S1). In normal skin, CIP2A staining was negative or weak in 60 of 67 cases (90%), while 7 of 67 cases (10%) showed strong staining (Figure 4F, Supplementary table S1).

### High CIP2A expression is associated with poor prognosis in cSCC

Kaplan-Meier survival analysis revealed high CIP2A expression was associated with poor disease-specific survival in the combined cohort of mcSCC and non-mcSCC patients (Figure 4G, log-rank p = 0.0410). This association was stronger when the analysis was restricted to patients with mcSCC (Figure 4H, log-rank = 0.0179). High CIP2A expression was also associated with worse overall survival among patients with mcSCC (Figure 4I, log-rank p = 0.0397). These findings indicate that high CIP2A expression is negatively associated with patient prognosis in cSCC.

## DISCUSSION

PP2A, is a serine/threonine phosphatase tumor suppressor that functions as a heterotrimeric complex composed of a scaffolding subunit, a catalytic subunit and one of several substrate-specific regulatory subunits (Nagelli and Westermarck 2024). PP2A is ubiquitously expressed in eukaryotic cells and inhibition of its activity in fibrosarcoma cells induces expression of invasion proteinase collagenase-1 (MMP-1) (Westermarck et al, 1994). PP2A activity is regulated by diverse mechanisms, including genetic mutations (Taylor et al 2019) and, primarily by interactions with endogenous inhibitors, such as the oncoproteins CIP2A, SET and PME-1, which are widely overexpressed in many human malignancies (Junttila et al 2007; Kauko and Westermarck 2018). CIP2A has been shown to be an important regulator of key cancer-related processes, including signaling cascade activity, mitosis and the DNA damage response (Nagelli and Westermarck 2024). Unlike CIP2A, SET and PME-1 are mainly located in the nucleus. Elevated SET levels have been detected in various cancer types, including both solid and hematological malignancies, and SET overexpression has been associated with chemoresistance and poor prognosis in breast cancer patients (Tozuka et al 2021). High PME-1 expression has been reported in several cancers and is associated with tumor cell growth. However, it has also been linked to improved prognosis in patients with rectal cancer, suggesting a dual role in cancer (Kaur and Westermarck 2016).

In this study, we investigated the role of PP2A inhibitors in non-melanoma skin cancer cSCC. Our results show that among the endogenous PP2A inhibitors, CIP2A expression was upregulated in cSCC cells, whereas expression of other PP2A inhibitors (SET and PME-1) did not differ between cancerous and normal cells as both were expressed abundantly in normal keratinocytes and cSCC cells. Because CIP2A was the only endogenous PP2A inhibitor studied that was upregulated in cSCC compared to normal skin, its role was investigated in more detail.

The expression of CIP2A *in vivo* was examined by IHC in a large panel of tissue samples representing normal skin and the progression from AK to invasive cSCC and cSCC metastases. CIP2A expression has previously been reported across a wide range of cancers, including breast, colon, ovarian and gastric cancers, as well as HNSCC (Junttila et al 2007; Côme et al 2009; Khanna et al 2009). In this study, CIP2A expression was observed in cSCC tumor cells *in vivo*. CIP2A expression was significantly higher in cSCC compared with that in AKs, cSCCIS or normal skin, and significantly higher in metastases than in primary cSCC. Additionally, strong CIP2A staining was associated with poor prognosis in cSCC patients. In contrast, no strong staining was observed in AK, and only a small fraction of cSCCIS samples showed strong CIP2A staining. These findings suggest that CIP2A may serve as a marker for invasive cSCCs and cSCC metastases.

The findings of the present study demonstrate that CIP2A is expressed in a subset of normal skin samples, with approximately 10% showing strong CIP2A staining. CIP2A, together with the DNA damage response protein TOPBP1 (topoisomerase II-binding protein 1), has been identified as a central regulator of mitotic DNA repair, essential for maintaining genome stability when conventional repair pathways are inactive (Martin et al 2025; de Haan et al 2025). In UV-exposed skin, CIP2A may therefore support the proliferation of basal keratinocytes, thereby facilitating the entry of DNA damaged cells into mitosis and initiation of cSCC, as previously demonstrated in basal-like breast cancers (Laine et al 2021). Consistent with this, CIP2A has been associated with the survival of cancer cells following DNA damage (Laine et al 2021). Tumorigenesis in cSCC is driven by a high mutational burden, largely attributable to cumulative UV exposure, and cSCC exhibits one of the highest mutation rates among malignant tumors (South et al 2014). Early events in cSCC pathogenesis include mutational inactivation of the tumor suppressor genes *TP53* and *NOTCH1* in epidermal keratinocytes (Piipponen et al 2021; Cho et al 2018; Pickering et al 2014). Additional commonly mutated genes in cSCC include *HRAS*, *CDKN2A*, *TERT* and *BRCA2* genes (Pickering et al 2014; South et al 2014; Al-Rohil et al 2016). Interestingly, inactivating *TP53* mutations have been reported to correlate positively with *CIP2A* expression via the p21–E2F1 axis (Laine et al 2013; Khanna and Pimanda 2016). These observations support a role for CIP2A as an important regulator of cSCC progression, potentially facilitating the entry of DNA-damaged cSCC cells into mitosis. Moreover, CIP2A is essential for the viability of mitotic cells deficient in homologous recombination (HR)-mediated DNA repair, such as those harboring BRCA mutations (Adam et al 2021). The CIP2A–TopBP1 complex also plays a critical role in clustering fragmented chromosomes during mitosis (Nagelli and Westermarck, 2024). Therefore, in cSCC, CIP2A might have functions beyond the regulation of PP2A activity, contributing to tumor cell survival through modulation of the DNA damage response.

The functional role of CIP2A was investigated in cSCC. Previous studies have demonstrated an important role for CIP2A in tumor growth. Inhibition of CIP2A expression with specific siRNA significantly reduced the growth of several xenograft tumors, including HNSCC, non-small cell lung carcinoma, and urothelial carcinoma (Junttila et al 2007, Xue et al 2013, Ma et al 2011). In this study, knockdown of CIP2A with specific siRNAs led to a significant decrease in cSCC cell growth and viability. The molecular mechanisms underlying CIP2A function in tumor growth are still under investigation. CIP2A has been shown to stabilize the oncoprotein Myc by inhibiting PP2A-mediated dephosphorylation of Myc in cancer cells (Junttila et al 2007, Niemelä et al 2012; Fang et al 2012). Moreover, Myc can increase CIP2A expression, establishing a positive feedback loop between Myc and CIP2A (Khanna et al 2009). In cSCC cells, CIP2A knockdown resulted in decreased phosphorylation of MEK1 and ERK1/2, key components of the MAP kinase signaling pathway implicated in cSCC progression.

Because high CIP2A expression is noted in about 70% of cancers and appears to nonessential for normal physiology, CIP2A is an attractive target for cancer therapy (Smith and Sheltzer 2022). CIP2A overexpression has been associated with poor therapy response in several cancers (Khanna and Pimanda 2016). Consistently, inhibition of CIP2A has been shown to sensitize cancer cells to a range of clinically used therapeutic agents. (Kauko et al 2020; Kauko et al 2018). Thus, tumor *CIP2A* status may serve as a biomarker of cancer aggressiveness and therapeutic sensitivity (Nagelli and Westermarck 2024). The molecular mechanisms underlying PP2A inhibition by CIP2A have been investigated to support the development of CIP2A-targeted inhibitors. Structural and functional studies have shown that amino acid substitutions within the CIP2A head domain abolish *in vivo* tumor growth in breast cancer models by both preventing PP2A inhibition and promoting CIP2A degradation (Pavic et al 2023). Drug-induced degradation of CIP2A would be particularly advantageous, as it would not only block CIP2A-mediated regulation of PP2A but also abolish additional functions of CIP2A, such as its direct interaction with TopBP1 (Pavic et al 2023). Although protein–protein interactions are traditionally challenging to target, recent advances support direct modulation of CIP2A-PP2A interactions using small molecules or peptidomimetics (Khanna et al 2013). In addition, RNA interference represents a potentially more effective strategy for targeting CIP2A, by inducing degradation of the entire CIP2A protein and thereby blocking its multiple functional roles (Khanna et al 2013).

The results of this study reveal that CIP2A is abundantly overexpressed in the invasive primary tumors and metastases of cSCC. Additionally, high CIP2A expression is associated with poor prognosis in patients with cSCC. In conclusion, these findings support CIP2A as an oncogene in cSCC and suggest that its overexpression is an important event in cSCC progression, promoting the growth of these non-melanoma skin cancers. The findings identify CIP2A as a biomarker and a potential therapeutic target in locally advanced and metastatic cSCCs.

## Author Contribution

Conceptualization: LN, VMK, PR; Funding acquisition: LN, VMK, PR; Investigation: LN, PT, LH, JH, JSK, PR; Methodology: LN, PT, LH, PR; Data curation: LN, PT, LH, PR; Supervision: LN, PR, VMK; Visualization: LN, PT, LH, PR; Writing—original draft preparation: LN, PT, LH, JW, VMK, PR; Writing—review and editing: LN, PT, LH, JH, JSK, JW, VMK, PR. All authors have read the manuscript and approved the submission.

## Declaration of generative AI and AI-assisted technologies in the writing process

During the preparation of this work the authors used OpenAI-5.4 in order to revise the language during the final editing the manuscript. After using this tool, the authors reviewed and edited the content as needed and take full responsibility for the content of the publication.

## Acknowledgements

We thank Johanna Markola and Sinikka Collanus for expert technical assistance and Dr. Reidar Grénman for cSCC cell lines. We thank the Histology Core of the Institute of Biomedicine at the University of Turku and the Bioinformatics Unit of Turku Bioscience. The Unit is supported by University of Turku, Åbo Akademi University, and Biocenter Finland. This study was supported by the Research Council of Finland (Grant 363211) Sigrid Jusélius Foundation, the Cancer Foundation Finland and the State Research Funding of the Turku University Hospital (Grant 13336) and by personal grants from the Cancer Foundation of Southwest Finland (LN and PR); Finnish Dermatological Society (PR); Finnish Cultural Foundation (PR); Finnish Medical Foundation (PR) and Turku University Foundation (LN).

## Abbreviations

PP2A: protein phosphatase 2A;
CIP2A: cancerous inhibitor of PP2A;
SET: phosphatase 2A inhibitor I2PP2A;
PME-1: protein phosphatase methylesterase-1;
RNA-seq: RNA-sequencing;
cSCC: cutaneous squamous cell carcinoma;
cSCCIS: cSCC *in situ*;
AK: actinic keratosis;
IHC: immunohistochemistry;
qRT-PCR: quantitative real-time PCR;
UV: ultraviolet;
TMA: tissue microarray

