## Supplement information for "CIP2A is a biomarker for progression and poor prognosis in cutaneous squamous cell carcinoma"

### CIP2A is a prognostic biomarker and therapeutic target in cutaneous squamous cell carcinoma

#### Supplementary Figures S1-S4

#### Supplementary Table S1

#### Whole western blots

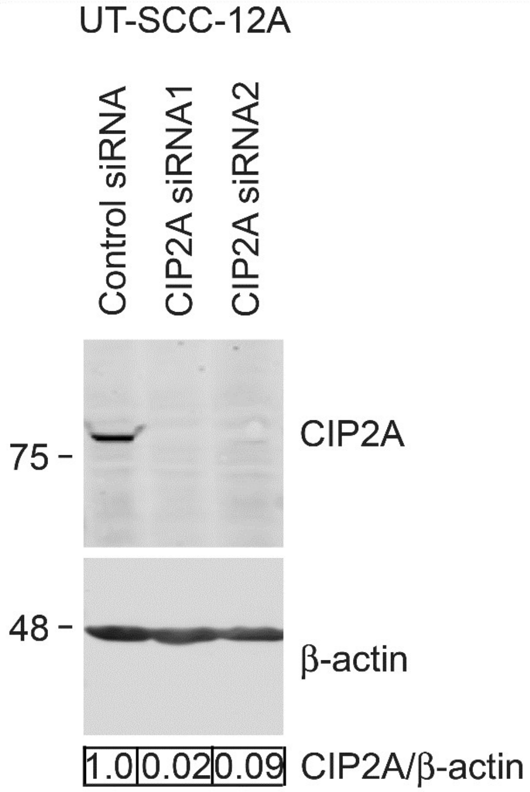


**Supplementary Figure S1. Knockdown of CIP2A with specific siRNAs.** Cutaneous SCC cells (UT-SCC-12A) were transfected with 75 nM negative control (Control siRNA) or CIP2A siRNAs. CIP2A level was determined with western blotting after 72 h. β-actin was used as a loading control. Migration positions of molecular weight markers (in kDa) are shown on the left.

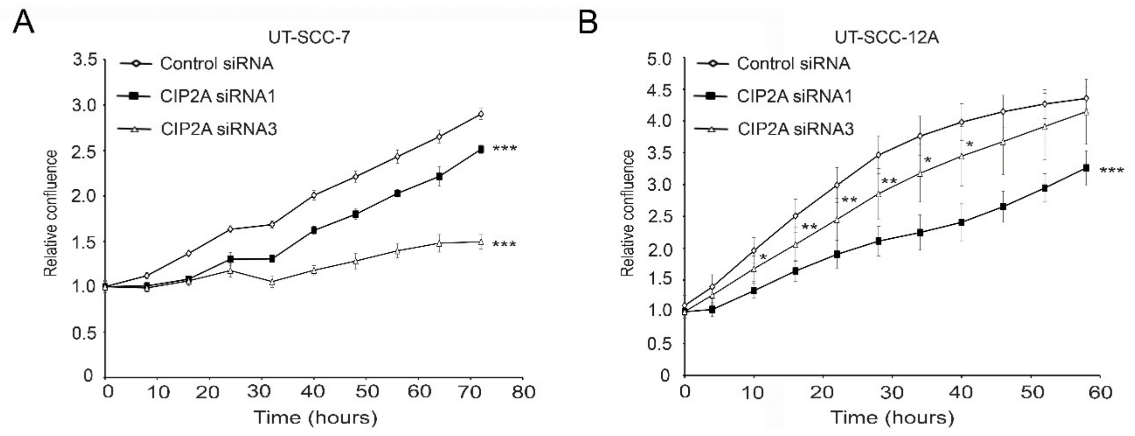


**Supplementary Figure S2. CIP2A regulates the growth of cSCC cells.** Cutaneous SCC cell lines (A) UT-SCC-7 and (B) UT-SCC-12A were transfected with 75 nM negative control (Control siRNA) or CIP2A siRNAs. The confluency of cSCC cells after CIP2A siRNA transfections was determined using IncuCyte S3 (UT-SCC-7 n=8; UT-SCC-12A n=6-8). Two-tailed Student's t-test. *p<0.05, **p < 0.01, ***p < 0.001.

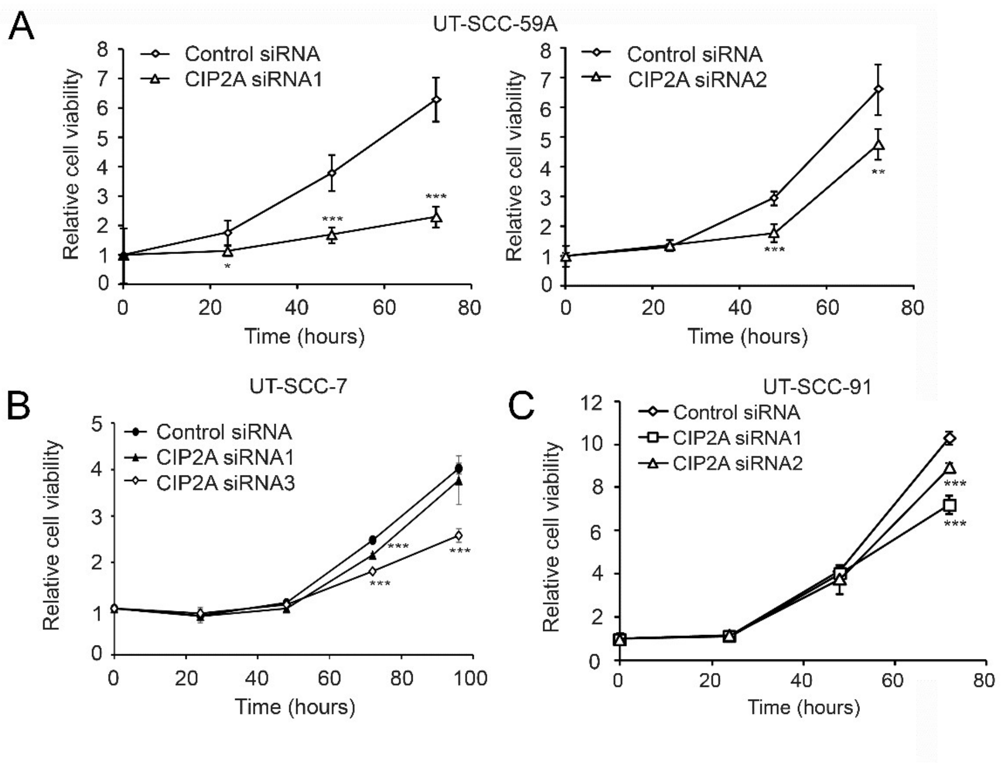


**Supplementary Figure S3. CIP2A regulates the viability of cSCC cells.** Cutaneous SCC cell lines (A) UT-SCC-59A, (B) UT-SCC-7 and (C) UT-SCC-91 were transfected with 75 nM negative control (Control siRNA) or CIP2A siRNAs. The number of viable cSCC cells was determined after siRNA transfection using CCK-8 assay at timepoints indicated (UT-SCC-59A and UT-SCC-7 n=8; UT-SCC-91 n=6-8). Two-tailed Student's t-test. **p < 0.01, ***p < 0.001.

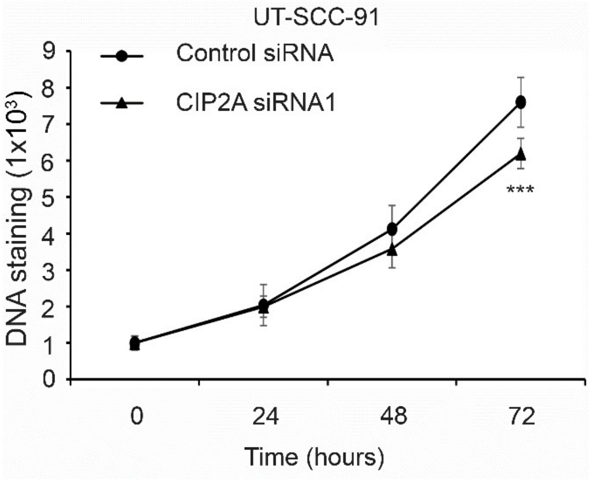


**Supplementary Figure S4. CIP2A regulates the number of cSCC cells.** Cutaneous SCC cells (UT-SCC-91) were transfected with 75 nM negative control (Control siRNA) or CIP2A siRNAs. DNA staining was performed 24, 48 and 72 h after siRNA transfection with SYTO 60 reagent and analyzed with LI-COR Odyssey CLx imaging system (n=7-8). Two-tailed Student's t-test. ***$p < 0.001$.

**Supplementary Table S1.** Increased staining intensity for CIP2A in tissue samples. Semiquantitative CIP2A staining intensity analysis. Number of specimens staining positive for CIP2A by IHC analysis in premalignant actinic keratosis (AK), cSCC *in situ* (cSCCIS), invasive cSCC and cSCC metastasis.

| Tissue sample | negative/weak staining | strong staining |
|---|---|---|
| Normal skin (n=67) | 60 | 7 |
| AK (n=73) | 73 | 0 |
| cSCCIS (n=59) | 58 | 1 |
| cSCC (n=162) | 80 | 82 |
| cSCC metastasis (n=63) | 12 | 51 |

**Whole western blots**

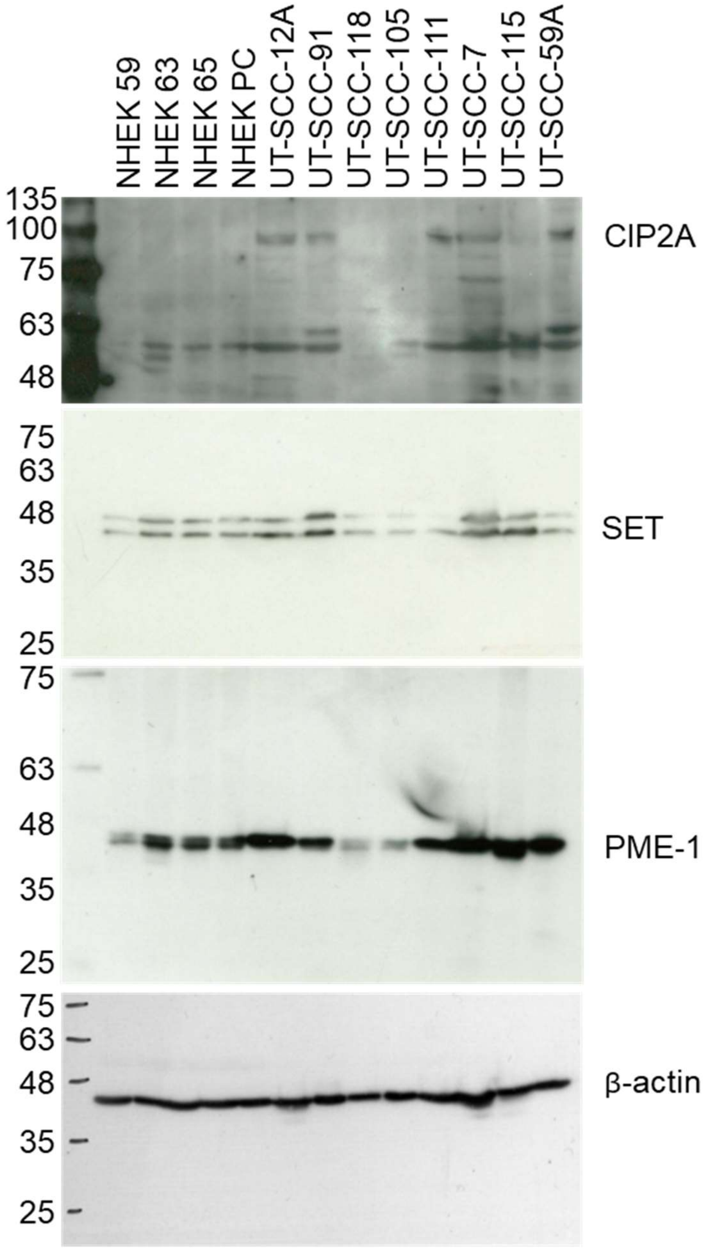


Figure 1D

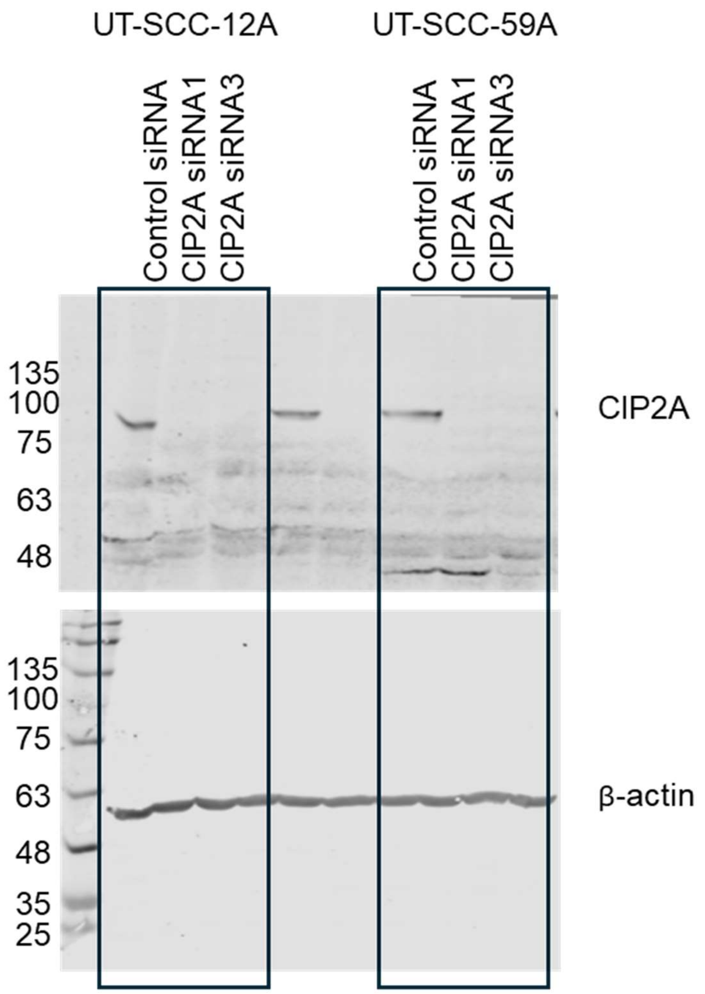
UT-SCC-12A
UT-SCC-59A
Control siRNA
CIP2A siRNA1
CIP2A siRNA3
Control siRNA
CIP2A siRNA1
CIP2A siRNA3
135
100
75
63
48
CIP2A
135
100
75
63
48
35
25
β-actin

Figure 3A

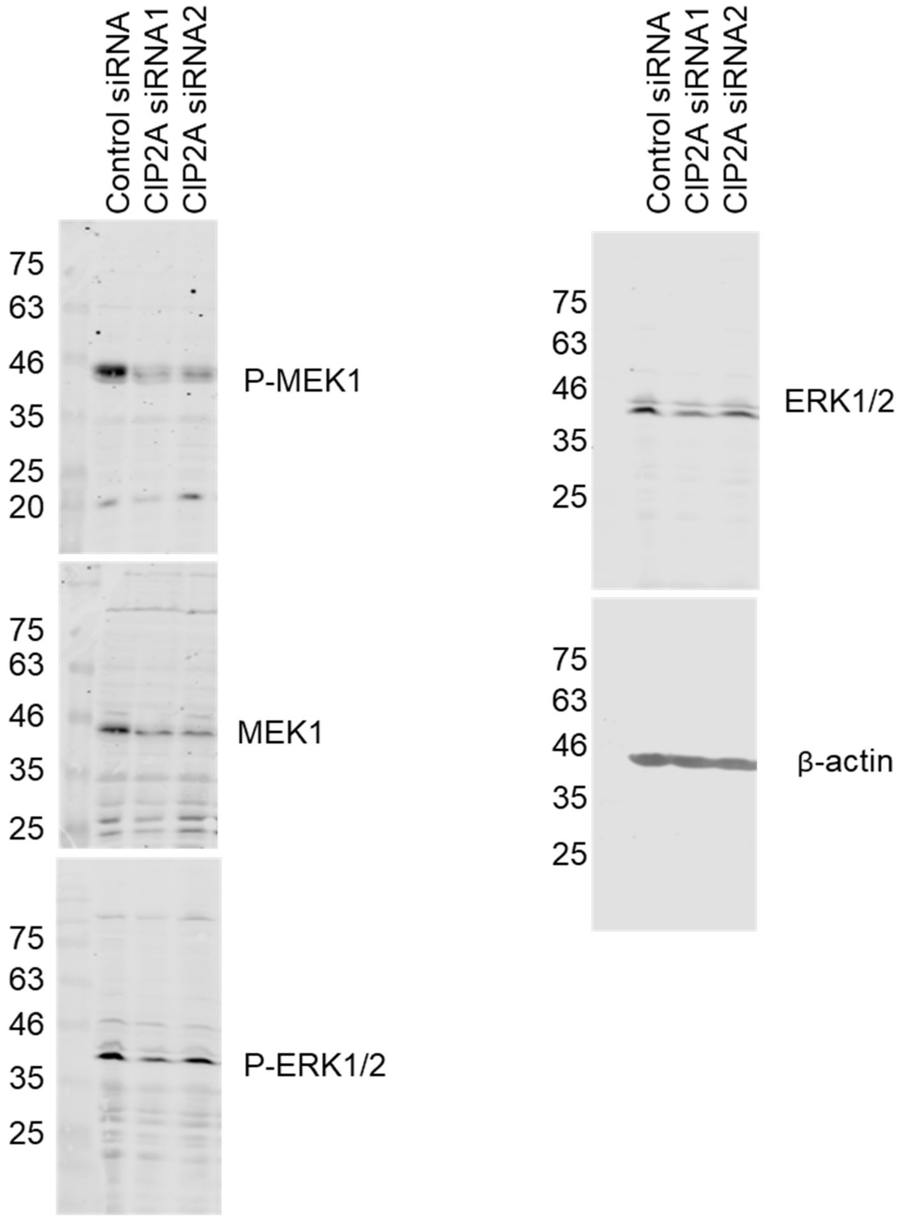
Control siRNA
CIP2A siRNA1
CIP2A siRNA2
75
63
46
35
25
20
P-MEK1
MEK1
P-ERK1/2
ERK1/2
β-actin

Figure 3E

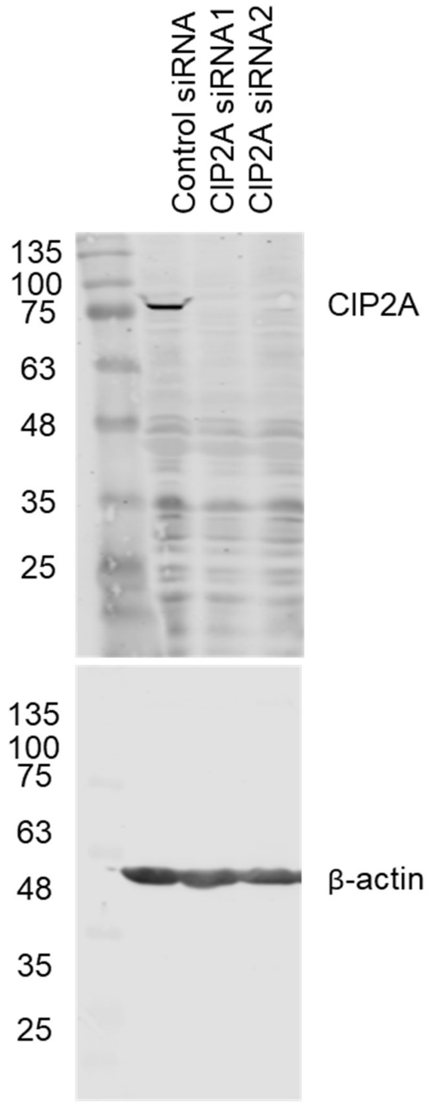


Supplementary Figure S1
